# Polyethylene glycol (PEG) methods are superior to acidification for secondary concentration of Adenovirus and MS2 in water

**DOI:** 10.1101/2021.11.19.469352

**Authors:** N.L. McLellan, S.C. Weir, H. Lee, M.B. Habash

## Abstract

Enteric viruses are a leading cause of waterborne illness worldwide and surveillance studies lack standardization in method selection. The most common and cost-effective approach to concentrating viruses from water samples involves virus adsorption and elution (VIRADEL) procedures, followed by secondary concentration. There is a lack of consistency in how secondary concentration methods are practiced and some methods may have better recovery for particular groups of viruses. Secondary concentration methods typically involve precipitation and the most common methods employ organic flocculation (OF) by acidification at a pH of 3.5, or precipitation by polyethylene glycol (PEG) in combination with the addition of NaCl. In this study, the recovery of coliphage MS2 using the plaque assay and human adenovirus strain 41 (HAdV41) using cell-culture and qPCR assays were evaluated by OF and PEG secondary concentration of spiked samples of wastewater, surface water, and groundwater. The recovery of MS2 and HAdV41 by PEG precipitation was significantly higher than that by OF (*p*<0.0001) when viruses were detected by culture based methods and marginally better when HAdV41 was enumerated by qPCR (*p*<0.019). The recovery of HAdV41 by qPCR ranged from 75.3% to 94.4% (*n*=36). The mean recovery of MS2 by OF was 4.4% (0.9%-7.7%; *n*=14) and ranged from 57.1% to 87.9% (*n*=28) for the PEG methods. The poor recovery of MS2 by OF was attributed to inactivation or poor stability at acidic conditions as MS2 were not recovered in the supernatant following OF and centrifugation. The inconsistency and lack of justification for method selection in many studies calls for a systematic study to inform guidance and standardization with respect to the application of concentration methods for various water types and viral pathogens.

**IMPORTANCE:** MS2 should not be used as a process control for methods involving acidification and culture-based detection. The dense floc produced by the PEG method may have contributed to higher recoveries as the pellet was more compact and stable than the loose pellet formed by OF. Standard methods for the detection of enteric viruses and surrogates that involve acidification could be modified with PEG precipitation to uphold virus recovery and minimize inactivation.

## INTRODUCTION

The detection of viruses in environmental waters (e.g. groundwater, surface water, and wastewater) typically requires isolation and concentration methods due to low titers and methodological sensitivity of detection methods. Nevertheless, low concentrations of viruses can pose a human health risk as the infectious dose of many viruses is low (e.g. 1 to 10 virions) (1, 2). Standard methods for the detection of enteric viruses in the environment are limited and many are not adaptable for all human viruses of concern or viral surrogates (e.g. phage); yet they are required to inform public health protection measures (3).

Procedures for detecting waterborne viruses typically begin with isolating viruses from the bulk water (e.g. groundwater or surface water) either by size-exclusion with ultrafiltration or with VIRADEL (virus adsorption-elution) methods (4–6). VIRADEL methods are most commonly employed due to costs and access to instrumentation. The limitations associated with these isolation and “primary concentration” methods for viruses from water have been reviewed elsewhere (5). The eluted suspension from a charged ultrafiltration and VIRADEL methods often requires secondary concentration prior to employing a culture- or molecular-based method for viral detection and quantification. Secondary concentration may be performed by a variety of methods, though the most common methods include organic flocculation (OF) by acidification and precipitation by polyethylene glycol (PEG) (5–8). Some matrices have elevated concentrations of viruses such as wastewaters, and these do not typically require the initial isolation step. Higher-titer samples can be concentrated directly by a “secondary concentration” method.

There is a lack of consistency in the application of secondary concentration methods with respect to: (a) the type of method selected and (b) the execution of each type of method. For example, PEG is often applied in combination with various molar concentrations of NaCl ranging from 0.2 to 1.5 M, and the incubation period may range from less than 1 h up to 24 h (7–13). The USEPA standard Method 1615 for the detection of enterovirus and norovirus in water employs OF by acidification for secondary concentration and recommends the use of poliovirus (Sabin poliovirus 3) as the process control virus (PCV) (6). The standard procedure for OF requires acidification of the suspension from an initial pH of about 9.0 (of the buffered beef extract used for elution of a cartridge filter) down to pH 3.5 ±0.1. Poliovirus has been shown to be resistant to acidification and drastic changes in pH. Huang et al. (2000) observed that poliovirus 1 was not inactivated by pH changes between 3.5 and 9.5 (14). The resistance of poliovirus raises the question as to its suitability as a PCV for virus detection methods; particularly if the methods are adapted for the detection of other, less stable, viruses.

Human adenoviruses (HAdV) have been proposed as a suitable surrogate for determining viral contamination in source waters due to their stability in environmental waters and persistence (5, 15–20). However, standard methods for the detection of HAdV are lacking. Further, the F-specific coliphage MS2 is commonly employed as a viral surrogate for understanding the fate and transport of viruses in the environment and for performance demonstrations of water treatment processes (e.g. ultraviolet light disinfection) (21–25). However, studies have indicated lower stability of some viruses during pH changes and in acidic environments (e.g. pH < 4), including MS2 (26–29).

The present study describes the recovery of MS2 and a human strain of adenovirus (HAdV41) from surface water, groundwater and wastewater by two secondary concentration methods: OF and PEG precipitation. Additionally, two molar concentrations of NaCl were trialed with PEG precipitation as varying concentrations have been cited in previous studies and to evaluate if this parameter is associated with improved recovery and resulting detection of viruses of interest (5, 8, 12, 14).

## RESULTS

### Background Water Quality

Raw groundwater was collected from a municipal well deemed as groundwater under the direct influence (GUDI) of surface water and had a turbidity of 0.16 NTU, temperature of 2.1°C, conductivity of 712 μS/cm, and absent of *E. coli* and total coliform detections at the time of collection. The raw surface water sample was collected in winter from the Grand River watershed which is heavily impacted (agriculture and urban land uses) and had a turbidity of 10.1 NTU, temperature of 4.2°C, conductivity of 577.13 μS/cm, *E. coli* concentration of 1.9×10^3^ CFU/100 ml, total coliform concentration of 9.1×10^4^ CFU/100 ml at the time of collection (30). Raw wastewater was collected from a municipal supply. Conductivity of the prepared 1L water samples of wastewater, concentrated surface water, and concentrated groundwater were 117, 61, and 124 μS/cm, respectively. Background concentrations of MS2 in the prepared 1L samples of wastewater, concentrated surface water, and concentrated groundwater were 5.6 log PFU/L (±5.3 log PFU/L), 4.1 log PFU/L (±4.1 log PFU/L), and 1.3 log PFU/L (±1.0 log PFU/L) where *n*=3 for each enumeration. Therefore, the spiked MS2 concentration target was >6 log MS2 PFU per L. Background concentrations of HAdV41 in the prepared 1L samples of wastewater, concentrated surface water, and concentrated groundwater were 4.6 log HAdV41, 2.0 log HAdV41, and 3.2 log HAdV41 gc/L, respectively. Therefore, the spiked HAdV41 concentration target was >5 log gc/L HAdV41, which is representative of concentrations found in the environment (31).

Spiked water samples contained 6.4 log MS2 PFU/L (±0.21 log MS2 PFU/L; *n*=9), 6.7 log HAdV41 IU/L (±0.75 log HAdV41 IU/L; *n*=6), and 7.7 log HAdV41 gc/L (±0.78 log HAdV41 gc/L; *n*=9); and these values were used as the baseline to determine virus recovery.

### Floc and Pellet Formation

The visual appearance of the floc in each suspension was different (**FIG**). Following OF, surface water samples did not have visible floc and the suspensions appeared clear (**FIG A**). Floc was more visible in the surface water suspensions flocculated by PEG (**FIG B**). Based on qualitative observations, the larger flocs formed by the PEG methods produced pellets which were larger, more visible, and more dense and opaque in the centrifugation vial when compared to the flocs formed by the OF method. These flocs were more robust; they remained intact while removing the supernatant. No difference was observed in the floc formation between the two NaCl concentrations used for the PEG methods.

### Recovery of Viruses

The recovery of MS2 and HAdV41 by OF and PEG methods from groundwater, surface water, and wastewater samples are presented in **FIG**. MS2 was evaluated by a culture-based method as this is the typical approach for the quantification of phage; while HAdV41 was detected by cell culture (TCID_50_) and qPCR.

#### (i) Recovery of MS2

The mean % recovery of MS2 PFU by OF was 4.3% (range of 1.0% to 7.6%; *n*=12) and was between 61 and 90% by the PEG methods. The recovery of MS2 by OF was significantly lower than that of the PEG method with either salt concentration (*p*<0.0001). There was no significant difference found between the recovery of MS2 by the two PEG methods (*p*=0.976). There was no significant difference between the recovery of MS2 from the various water types (*p*≥0.210). While it may appear that the recovery by the PEG methods is more variable than that of the OF method, the low recovery of OF may bias this observation.

#### (ii) Recovery of HAdV41 enumerated by cell culture

There was a high degree of variation in the recovery of HAdV41 infectious units (IU) between replicates, concentration methods, and water type (**FIG B**). The recovery of HAdV41 IU by PEG methods performed on groundwater samples (range 19.5-51.3%; *n*=6) was lower than that achieved for surface water (range 55.7-100.4%; *n*=6) and wastewater (range 71.8-102.2%; *n*=6) samples. The recovery of HAdV41 IU by OF for all water types ranged from 6.9-50.7% (*n*=9). For groundwater samples, there was no significant difference between the recovery of OF and PEG methods for HAdV41 IU (*p*>0.674). There was no significant difference between the recoveries of the PEG methods with different NaCl concentrations for any water type (*p*=0.482). The recovery of HAdV41 IU was significantly higher by PEG with either salt concentration than the OF method for surface water and wastewater samples (*p*<0.0001).

#### (iii) Recovery of HAdV41 enumerated by qPCR

The recovery of HAdV41 enumerated by qPCR ranged from 75.3-94.4% by all secondary concentration methods for all water types (**FIG C**). The recovery for wastewater samples was significantly higher than that achieved for groundwater samples by both PEG methods (*p*<0.003); though the variability was substantially lower than when HAdV41 was detected by cell culture.

#### (iv) MS2 Recovery in Supernatants

MS2 is a common PCV and was enumerated in the supernatant of each test to evaluate the fate of MS2 that was not recovered in the pellet following secondary concentration by OF and PEG methods. The spiked concentration of MS2 was used as a baseline to calculate the average recovery of MS2 in the pellet, supernatant, and the resulting “unaccounted” fraction which may indicate the inactivated MS2 not detected by the plaque assay; the results are shown in **Fig. 3**. The recovered fraction of MS2 in the supernatant ranged from 1.1% to 20% for the PEG methods, and from 13.9% to 34.8% for the OF method tested. The unaccounted fraction of MS2 from the PEG methods ranged from 5.9% to 29.1% for the PEG methods, and from 58.7% to 80.9% for the OF method tested. The recovery of MS2 in the supernatant was lowest, and the fraction of unaccounted MS2 was highest, in surface water samples for all secondary concentration methods tested.

**FIG 1.**
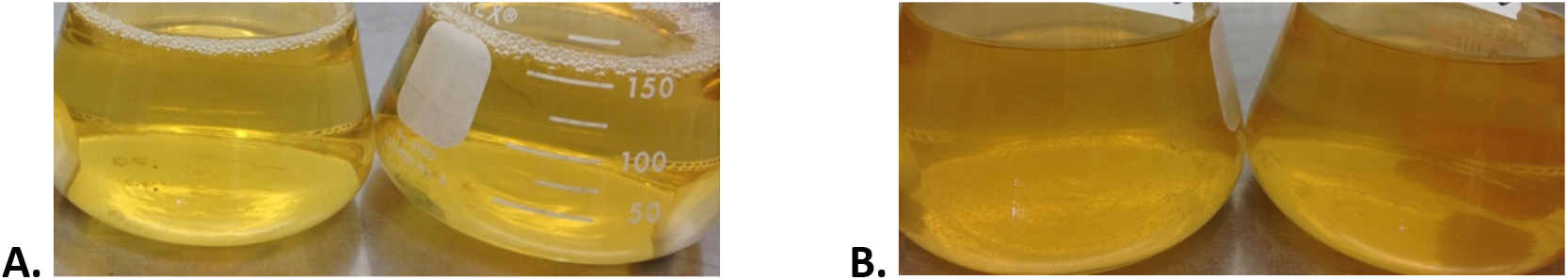
Flocculated suspensions of surface water. Duplicate suspensions following OF lacking visible flock (**A**); Suspensions following flocculation using 0.5 M PEG (left) and 1.5 M PEG (right) with visible floc (**B**).

**FIG 2.**
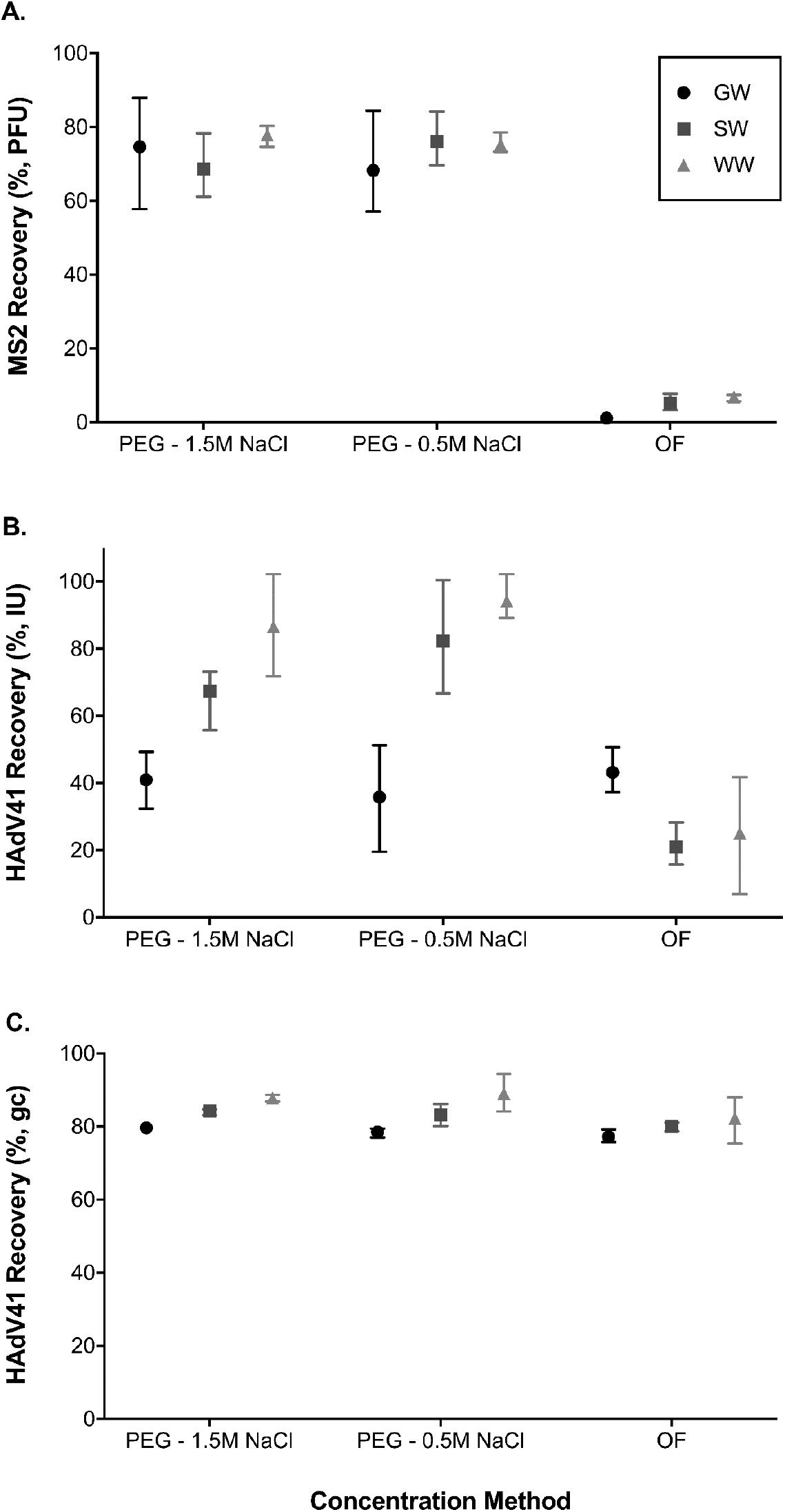
Recovery of MS2 in percent plaque forming units (PFU) (**A**), HAdV41 enumerated by cell culture in percent infectious units (IU) (**B**), and HAdV41 enumerated by qPCR in percent gene copies (%) (**C**), by three concentration methods (PEG with 1.5 or 0.5 M NaCl and organic flocculation [OF]) performed on wastewater (WW, 1L) and BE NanoCeram^®^ elution from groundwater (GW; 1000 L) and surface water (SW, 80 L). Error bars show the range of results (*n*=4 for each treatment).

**FIG 3.**
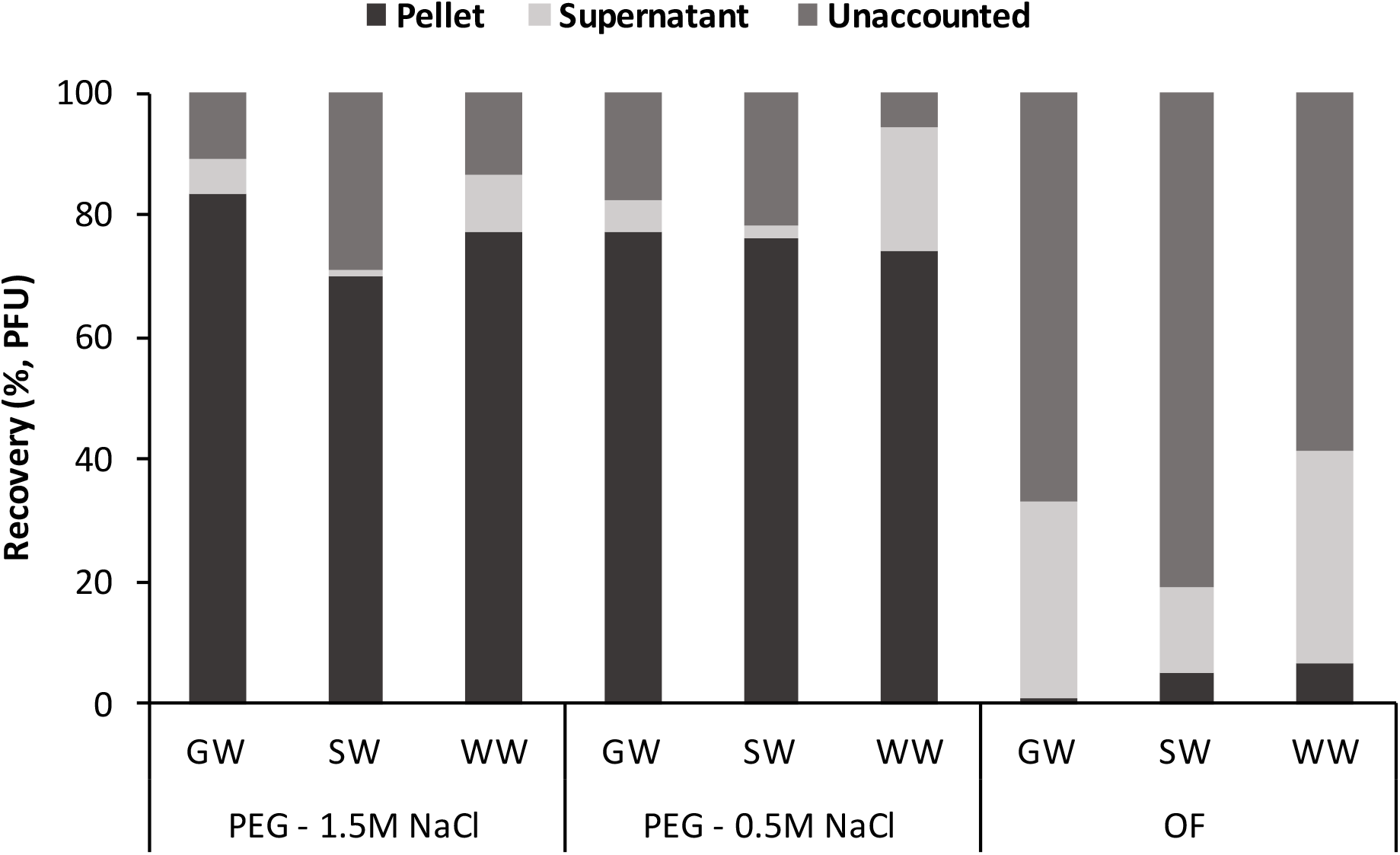
Average recovery of MS2 PFU in the supernatant, and in the suspended pellet following flocculation, by the PEG method employing two concentrations of NaCl and OF for three water matrices; groundwater (GW), surface water (SW), and wastewater (WW), and the resulting unaccounted fraction of MS2 based on a mass balance using the initial spiked suspension concentration as a baseline (*n*=4 for each test).

**FIG 4.**
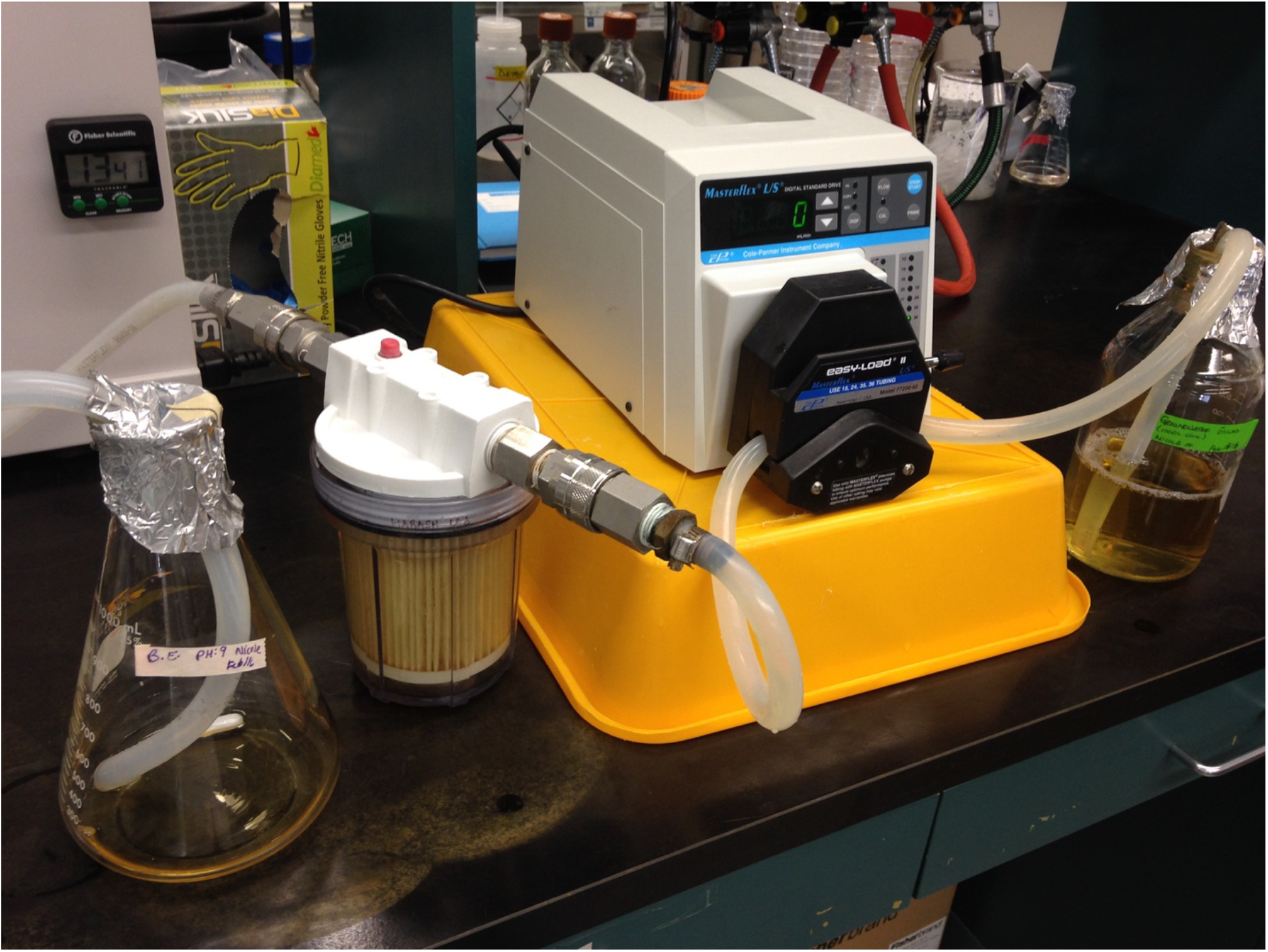
Virus adsorption and elution (VIRADEL) bench-top apparatus using NanoCeram^®^ positively charged pleated cartridge filters.

**FIG 5.**
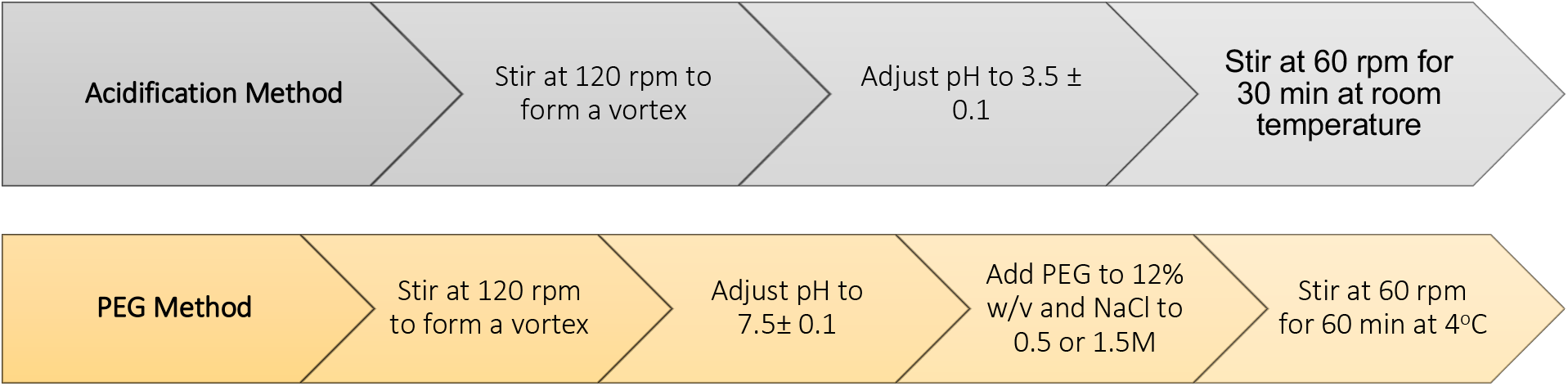
General steps for acidification and PEG flocculation methods.

**FIG 6.**
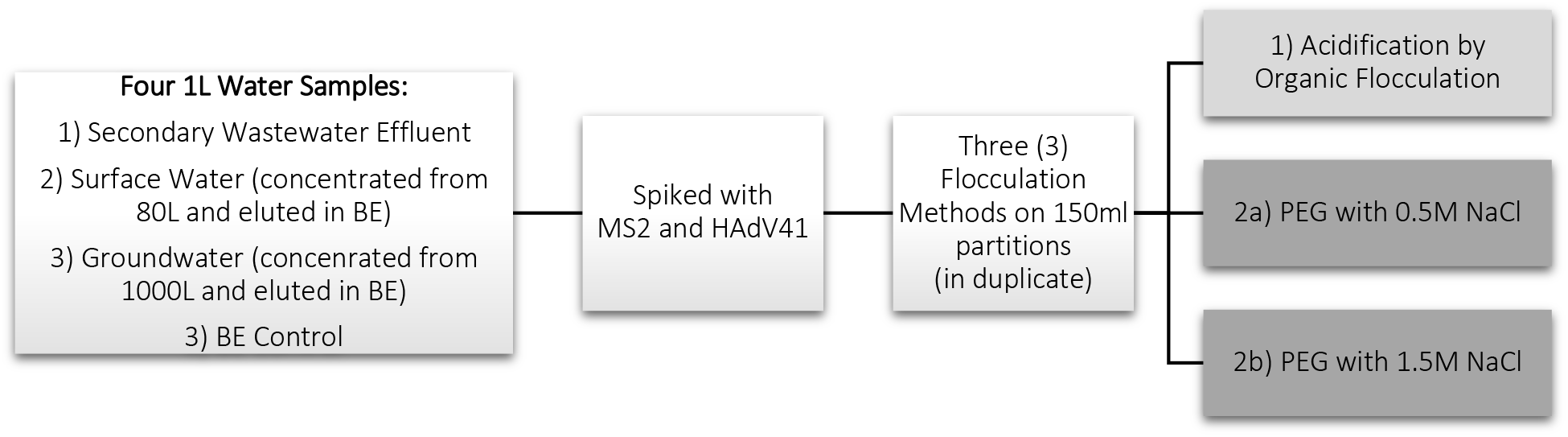
Sample preparation and partitioning for three flocculation methods: 1) acidification and 2) two PEG methods. The procedure was performed with duplicate samples and repeated to produce *n*=4.

## DISCUSSION

In this study, the recovery of MS2 and HAdV4 by PEG precipitation, when detected by culture-based tests, was found to be superior to that of OF by acidification. The >90% loss of culturable MS2 by OF is suspected to be primarily due to the inactivation of MS2, which is known to have poor stability in acidic environments. Acidification has been reported to impact virus integrity and infectivity by others (32, 33). Langlet et al. (2007) observed a mean 3 log (±0.99; *n*=12) decrease in MS2 concentration at pH 2.5 when the phase was stirred for 4 h in buffered KCl solution (0.4 M) at 20°C, as compared to a loss of 1.11 log (±0.39; *n*=5) at pH 3.9 and an insignificant change of +0.23 log MS2 (±0.15; *n*=8) at pH 6.7 (29). A larger fraction of MS2 was recovered in the supernatant of samples that were concentrated by OF than by PEG methods which may reflect the greater stability of floc formed by PEG precipitation. Additionally, a substantially larger fraction of MS2 was unaccounted for from OF concentration than from PEG, which may indicate greater inactivation of MS2 during sample processing during the OF method. Thus, MS2 should not be used as a process control virus (PCV) for methods involving acidification when culture-based detection is used. Further, PEG precipitation outperformed OF in terms of the recovery of adenovirus detected by cell culture in surface water and wastewater samples, and had similar recovery to OF in the groundwater samples.

The concentration of viruses from liquid suspensions using PEG precipitation began prior to the 1970’s and has been evaluated for the concentration and recovery of a suite of enteric viruses. Yamamoto and Alberts (1970) experimented with the removal of a variety of bacteriophage types using 2-10% solutions of PEG_6000_ (34). Thereafter, Lewis and Metcalf (1988) found that PEG precipitation was more effective than OF for the concentration of rotaviruses (WA and SAII) and hepatitis A virus (HAV) from estuarine and fresh waters (11). PEG_6000_ was added to each sample (8%) and stirred for 1.5 h at 4°C, followed by centrifugation at 10,000 *g* for 20 min. The pellet was reconstituted in phosphate buffer solution (PBS). Various concentrations of PEG_6000_ were tested (0, 8%, 12%, 15%, 20% w/v), and between 12-15% w/v was found to be optimal for viral suspensions. Nevertheless, some studies continue to perform PEG methods with PEG concentrations lower than 12% w/v (9, 28), and others have found that higher PEG concentrations can enhance the recovery of some viruses (e.g. TGEV and HAV at 20% w/v PEG) (12, 35).

The SARS-COV-2 pandemic has sparked a renewed interest in evaluating, optimizing, and standardizing methods for the concentration of viruses in wastewater in an effort to improve wastewater-based epidemiology (WBE). Peer-reviewed studies have compared PEG precipitation with other concentration methods for the recovery of a range of viruses and relevant surrogates (36–40). PEG precipitation (10%) has been preferred by a number of studies for providing the highest recovery of viruses and surrogates, and for its simple approach (i.e., low-cost equipment, accessible, no pre-treatment steps) (41–46). Many of these recent findings are aligned with the results of this study, and Lu et al. (2020) found that PEG methods were among the most common due to reliability, broad applicability, and accessibility (45). Pecson et al. (2021) demonstrated that PEG methods, particularly those excluding solids removal, were the most reproducible and produced the highest recovery of gene copies for SARS-COV-2 from raw wastewater samples when compared to ultrafiltration and acidification methods (40). Ahmed et al. (2020) found that methods that included acidification to pH 4 produced the poorest recovery of murine hepatitis virus (47). Nour et al. (2021) found that a PEG precipitation method (10% w/v) produced the highest recovery of HAdV DNA from treated wastewater using a factorial comparison approach against glass wool-based concentration and charged membrane based adsorption/elution (44). Torii et al. (2021) compared PEG precipitation for the recovery of *Pseudomonas* phage ∂6 (as a surrogate for enveloped viruses) with that of non-enveloped MS2 by RT-qPCR and found that MS2 showed differences in recovery and was not a suitable indicator to validate the extraction and recovery of enveloped viruses (42). This recent interest in evaluating various concentration methods has highlighted the inconsistencies in how PEG methods are applied and the lack of standardization; as many of these studies include or exclude pre-treatment steps, use a range of PEG concentrations (e.g., up to 20% PEG (48)), and a range of incubation times from 4 h to “overnight”.

This study used a higher concentration of PEG (12% w/v) than has been used in other studies (5, 8). The recovery of viruses in this study are similar to, or greater than, those found by others that used PEG concentrations of 10% or less (10, 28). Enriquez and Gerba (1995) found no significant difference between the average recovery of HAdV40 (detected by cell culture) in tap water, sea water, and wastewater (secondary sewage) by OF (38.6%) and PEG precipitation (40%), where the PEG methods was performed with 7% PEG_8000_ and 0.5 M NaCl incubated for 2 h at 4°C (10). El-Senousy et al. (2013) found that PEG_8000_ (12%) with 1.5 M NaCl provided significantly better (average 0.4 log better; *p*<0.05) recovery than OF for norovirus GI and GII by RT-PCR from fresh produce and irrigation water (8). In another study, PEG was applied at 14% w/v with 0.2 M NaCl to wastewater samples with overnight incubation at 4°C, the recovery of poliovirus detected by plaque assay was within an acceptable range of 59.5% (±19.4%); though recovery of poliovirus was significant higher (106% ±1.6%) with an alternative method employing skimmed milk acidification which requires a pH of 3 to 4 (7). Ye et al. (2016) found the recovery of MS2 by PEG (8% w/v and 0.5 M NaCl) using a plaque assay was about 43.1% (±16.8%) during the concentration of municipal wastewater samples (28).

A combined approach, using acidification and PEG methods, has been used for secondary concentration during surveillance of enteric viruses in environmental waters (49). Farkas et al. (2018) acidified the beef extract elution from a VIRADEL primary concentration method to a pH of 5.5 and then held for 30 min followed by centrifugation to remove sample particulates (2,500 x*g*, 10 min). The supernatant received 15% w/v PEG_6000_ treatment with 2% NaCl with an overnight (16 h) incubation at 4°C followed by centrifugation (10,000 x*g* for 30 min, 4°C). This approach resulted in an average recovery of adenovirus gc by qPCR of about 60% from a river water source which was similar to the recovery of the process control virus mengovirus of 59.5% (40.6 standard deviation). The authors suggest that the performance of this method for a range of sample types (wastewaters, surface water, sediment, and shellfish samples) could be standardized for routine monitoring. However, the necessity for the acidification step is unclear and for many waters (lake and groundwater sources) the initial centrifugation step would not be of benefit; in fact, it could contribute to additional losses of particle-associated viruses. The recovery of HAdV41 gc was consistently >60% for all waters and secondary concentration methods used in this study; therefore, the use of a hybrid method is not recommended.

PEG methods have been performed with multiple washing steps of the pellet (50) which may negatively affect the recovery of viruses. While efforts should be made to minimize the co-concentration of inhibitory compounds to qPCR and cell-culture assays with target viruses (51), recovery should be upheld as a primary goal through viral concentration from environmental water samples even when a process control is included. When PCR methods are employed, the efficiency of nucleic acid extraction can be equally important to the concentration step, and a review of commercial extraction kits has been reported by Iker et al. (52).

Masclaux et al. (2013) used a direct PEG precipitation method (32% w/v PEG_8000_, 1.2 M NaCl without primary concentration) for the concentration of enteric viruses in wastewater and the average recovery efficiency for hepatitis E virus was 39% (ranged from 25-53%, *n*=5) and for the PCV Rice Yellow Motile Virus (RYMV) was an average recovery efficiency of 66% (ranged from 58-71%, *n*=5) when detected by PCR. Direct PEG treatment of waters with high viral titers and elevated solids is practical as it eliminates the pre-filtration and primary concentration steps which can have issues with filter clogging (12, 53, 54) or de-adsorption of viruses off of filter material (55). Further, environmental waters with high viral titers (e.g. >10^3^ IU/L), such as wastewaters and impacted surface waters, are likely to be the most commonly monitored aquatic sites for enteric viruses as discharge regulations become more stringent and watershed managers are driven to understand pathogen concentrations in their systems to inform risk assessments (3, 56, 57). Masclaux et al. (2013) centrifuged wastewater samples prior to performing direct PEG precipitation on the supernatant to minimize PCR inhibition from organics, though this step may have reduced the recovery efficiency of viruses associated with particles (12).

The dense floc produced by the PEG precipitation method in this study could contribute to higher recoveries and higher consistency between replicates as the pellet was more compact and stable than the pellet formed during OF. Further, PEG precipitation presents a more user-friendly method whereby the analyst can visually confirm that the suspension was effectively centrifuged during the pelleting procedure and that the pellet is intact during decanting the supernatant. A second centrifugation step (8000 x*g* for 5 min at 4°C) to further compact the pellet has been proposed to enhance virus recovery by flocculation methods, though the benefits to recovery from this additional step have not been confirmed (9). In this study, the supernatant was extracted by pipette, rather than by decanting or physical pouring. This approach allowed the centrifuge container to remain stable and caused minimal disturbance to the pellet.

The results from this study suggest that the recovery of HAdV41 nucleic acid may be improved for samples with higher turbidity or solids content as higher recoveries were achieved for the wastewater sample, followed by the surface water and groundwater sample; while the recovery of MS2 was not substantially impacted by water type. The particulate matter in more turbid samples may provide a *de facto* flocculant-aid that provides greater surface area or bridging to enhance the formation of larger flocs, and higher molecular weight for better sedimentation into the pellet during centrifugation. PEG methods have been found to be more commonly applied to surface water samples, while OF methods have been more commonly used to concentrate wastewater samples (58). This tendency may be the result of laboratory preferences or the fact that wastewater samples typically have higher viral titers and a loss of viruses during acidification may be negligible and allow for adequate detection. The inconsistency and lack of justification for method selection in many studies calls for a systematic study to inform guidance and standardization with respect to the application of concentration methods (3).

Fout and Cashdollar (2016) indicated that USEPA Method 1615 may be adapted for the detection of adenoviruses (59). It is recommended to explore modifying existing methods with secondary concentration by PEG precipitation to uphold virus recovery and minimize inactivation during sample processing and detection by culture-based methods. Additionally, PEG and OF methods should be compared for other viruses of interest to determine if there are significantly different performance characteristics. Standard methods for the detection of enteric viruses in the environment may require customization for virus families due to the diversity in virus stability, and water type. There is not likely one method that would produce a high recovery of all viruses and surrogates of interest as there is high phenotypic and biochemical diversity among viruses and phages. For example, all enteric viruses and F-specific phage are non-enveloped and considered to be more stable in the environment than enveloped viruses (e.g. influenza, coronaviruses, hepatitis B) (28, 35). While this information may add complexity to existing multi-step procedures, this guidance will benefit the quality of information that is gathered to inform risk assessments of recreational and drinking waters.

Appropriate process controls are necessary to provide an indication of the performance of virus detection methods and provide a level of quantification with respect to recovery. However, the recovery of a PCV does not provide an understanding of the mechanisms that contributed to virus losses such as inactivation, poor de-adsorption during elution, or poor flocculation unless the PCV has properties similar to the target virus. MS2 has been proposed and used in numerous studies as a PCV (5, 60–64). However, the results of this study suggest that the use of MS2, and other surrogates that may be unstable during sample processing, should be done with caution or avoided in the case of acidification protocols when culture-based detection is employed. Alternatively, murine norovirus (MNV1) has been suggested as a PCV (9, 22) and it has been found to be stable across the pH range from 2 to 10 (27).

A limitation of PEG methods is that the chemical conditions may contribute to the inactivation of enveloped viruses by disrupting the lipid bilayer during concentration as has been seen for influenza and murine hepatitis virus (+ssRNA, coronavirus), which may be attributed to the NaCl concentrations (28). The selection of a secondary concentration method may be affected by the limitations of the detection method. For example, where PCR detection methods are employed (e.g. for the detection of noroviruses which do not have established culture-based methods), the inactivation of viruses may not be a primary concern so long as no damage is caused to the genome during sample processing. However, secondary concentration methods that have improved virus recovery and reduced co-concentration of PCR-inhibitors would be beneficial for PCR detection methods. A perceived limitation of PEG may be the additional time to perform incubation. Typical PEG methods are performed with an overnight (i.e. 16 h) incubation at 4°C (though several studies report incubation times of 1 to 4 h), while OF methods are typically completed in less than 1h. While PEG methods may add time to the procedure, it does not necessarily add more labor as the incubation is unsupervised. The priorities of method selection should be to uphold virus stability and recovery of viruses to produce the most accurate quantification when it comes to an assessment of human health risk. An overnight incubation step still allows for these methods to provide results in a 1-day turnaround when coupled with qPCR. This study and several others have completed PEG methods with acceptable results with an incubation time of 1h.

In conclusion, this study presents primary data indicating the inactivation of coliphage by OF acidification during secondary concentration of VIRADEL detection methods; though the results from this study are applicable to elutions from ultrafiltration as well. Acidification methods for the concentration of viruses should only be employed where viral stability at low pH conditions has been confirmed or when only molecular based assays are used for virus detection. Other alternative methods, such as PEG precipitation, are available that can produce comparable or improved recoveries during concentration procedures that substantially minimize the inactivation of enteric viruses for culture-based detection.

## MATERIALS AND METHODS

### Virus Propagation and Enumeration

#### MS2 bacteriophage

MS2 bacteriophage was propagated according to methods previously described (65, 66). Briefly,1 ml of 10^8^ PFU/ml MS2 (ATCC #15597-B1) was inoculated into 800 ml tryptic soy broth (TSB) with log phase *Escherichia coli* C3000 cells (ATCC #15597) and shaken at 120 rpm for 4 h. Propagated MS2 was isolated by first centrifuging down the *E. coli* host at 8000 *g* for 10 min, followed by filtering the supernatant through a 0.45 μm pore membrane (sterile polyether sulfone filter, VWR North America, USA). Concentrated MS2 stocks were stored at 4°C for short durations (<2 days) or at −80°C for longer durations (17, 22, 67).

MS2 was enumerated using the single-layer agar method (68). Plates were incubated at 37°C for 36 to 48 h. Circular clearings in the agar were enumerated and recorded as plaque forming units (PFU) per ml of sample.

#### HAdV41 Propagation

HAdV41 is a non-enveloped double-stranded DNA virus. It was used in this study as a model human enteric virus as it is ubiquitous in environmental waters and has potential utility for microbial source tracking (15, 49, 69–73). HAdV41 was acquired from the laboratory of Prof. Martha Brown, Dept. Laboratory Medicine and Pathobiology, University of Toronto. HAdV41 was propagated and enumerated according Leung and Brown (2011) (74). HAdV41 was cultured using cell line HEK 293 (human embryonic kidney cells), at passages 40 to 80 in complete growth medium containing 350 ml 1X MEM (Corning™ cellgro™ Minimal Essential Medium Eagle Mod. with Glutamine), 40 ml fetal bovine serum (FBS) (Hycone™, GE Healthcare Life Sciences), 4 ml penicillin-streptomycin (10,000 U/ml Pen Strep, Gibco Life Technologies), and 4 ml 7.5% sodium bicarbonate (7.5%, Gibco), and adjusted to pH 7.4 ± 0.2 using 1M HCl.

#### HADv41 Detection by Culture

Infectious units (IU) of HAdV41 were enumerated by an endpoint dilution assay (TCID_50_) using 96-well clear polystyrene flat bottom microplates (8 rows by 12 columns) (Corning) conducted in duplicate. Each water sample was diluted on a separate plate. All interior wells received 90 μl of growth medium including the control row; exterior wells (around the entire perimeter of the plate) were filled with sterile water to inhibit evaporation. Each of the 10 wells in the second row were inoculated with 50 μl of HAdV41 sample, and a multi-channel pipette was used to serially dilute the sample down the subsequent five rows by transferring 10 μl and changing the tips after each row passage. One 25 cm^2^ flask with confluent growth of HEK 293 cells was used to add cells to each 96-well plate. Trypsinized cells were suspended in about 12 ml of culture medium to achieve a concentration of about 10^5^ cells/ml (determined using a hemocytometer), and about 200 μl of this cell suspension was added to each well containing a sample. Positive control wells were inoculated with the HAdV41 stock and negative control wells inoculated with sterile PBS were included in each plate. Plates were incubated (37°C, 5% CO_2_) until 10 d post infection (p.i.). Each well was examined at 40x magnification under an inverted light microscope. Evidence of cytopathic effect (CPE) was recorded as either present (+) or absent (−). The ratio of infected to uninfected wells was used to calculate the Log TCID_50_ (tissue culture infectious dose). With the assumption that 0.7 IU per well represents the 50% endpoint, the concentration of IU of HAdV41 in the original samples was calculated as IU/L.

#### HADv41 Detection by PCR

HAdV41 was enumerated by qPCR using the extraction method and primer probes described previously (75). Nucleic acid was extracted from 200 μl of each sample using the QIAamp MiniElute™ Virus Spin Kit nucleic acid (for DNA and RNA) according to the manufacturer (QIAgen Canada, Toronto, ON, Canada). DNA samples were stored at −80°C and assayed in duplicate by qPCR. Each qPCR assay was performed in 50 μl with PCR master mix (iQ™ Multiplex Powermix, Bio-Rad Life Science), 250 nM primers and 100 nM probe, and 10 μl of DNA extract. The thermocycler (iQ5 Multicolor Real-Time PCR Detection System, Bio-Rad) conditions included one cycle at 95°C for 10 min, 40 cycles of 95°C for 15 s followed by 60°C for 1 min. The standard curve was prepared in a series of 10-fold dilutions from 10^1^ to 10^8^ copies of plasmid/well. It was assumed that one copy number was the equivalent of one enteric virus unit. The efficiency of the qPCR assay was 101.5% +/− 5% with R^2^ values of 0.98-1.00, and slope between −3.21 and −3.50.

### PCR Inhibition

PCR inhibition was determined by spiking a concentrated HAdV41 DNA stock (6.4×10^4^ gc/μl) into twofold serial dilutions (no dilution, 1:2, 1:4, 1:8) of a nucleic acid extract of each water type as well as into reagent-grade water (Milli-Q, Water Purification System, Millipore, Ontario) to serve as a positive control and subject to PCR as described above. The PCR inhibition test determined that the following dilutions of samples minimized inhibition of the HAdV41 qPCR assay by <1 Ct when compared to the positive control for the following water types: 1:2 for all groundwater samples; 1:4 for the prepared and spiked surface water samples; 1:8 for prepared and spiked wastewater samples; and 1:2 for all suspended pellet samples. These dilutions were used for all respective HAdV-qPCR reactions.

### Prepared Water Samples

One litre samples of each of the following water types were prepared: wastewater (WW; secondary effluent), concentrated groundwater (GW; from 1000 L), and concentrated surface water (SW; from 80 L). Bulk concentrated groundwater and surface water (river water) samples were prepared once by filtration through NanoCeram^®^ positively charged pleated cartridge filters (Argonide, Sanford, Florida, USA) at a maximum flow rate of 10 L/min. The goal of this step was to mimic the background chemistry of typical water samples that are collected using common adsorption/elution methods which precede the concentration and detection of enteric viruses in source waters. NanoCeram^®^ filters are patented; the manufacturer states that they are composed of non-woven thermally-bonded microglass fibers, cellulose and nanoaluminia to create a positive charge on the filter surface with a rating of 0.2 μm. NanoCeram^®^ filters were eluted with 1 L of beef extract (BE) at pH 9 according to USEPA (6) (**FIG**). The eluted samples were then analyzed for background concentrations of coliphage enumerated by the MS2 detection method, and HAdV41 by cell culture and qPCR.

### Experimental Design

Each 1 L prepared water sample was spiked with MS2 bacteriophage and HAdV41 and split into 6 portions of 150 ml each (**FIG**). Therefore, loss of viruses during the initial concentration and isolation phase of NanoCeram filtration was not assessed as part of this study and would not bias the study outcomes with respect to evaluating OF or PEG flocculation. Three flocculation methods were performed in duplicate; one acidification method by OF, and two methods using PEG with 0.5 and 1.5 M NaCl, respectively. Following flocculation and prior to concentration by centrifugation, 10 ml samples were collected from each 150 ml flocculation vessel to determine potential loss of infectivity and detection of viruses due to each flocculation process.

### Flocculation by Acidification

Flocculation by acidification was performed according to the OF method described by the USEPA (6). Spiked samples (150 ml each) were stirred using a magnetic stir bar and plate at 120 rpm to form a vortex and pH was adjusted to 3.5 ± 0.1 by adding 1M HCl. Samples were stirred at 60 rpm for 30 min at room temperature (21°C ± 1°C) to facilitate floc formation (**FIG**).

### Flocculation by PEG

Flocculation by PEG was performed according to (8) at two different concentrations of NaCl (**FIG**). Spiked samples (150 ml each) were stirred at 120 rpm to form a vortex and pH was adjusted to 7.5 ± 0.1 by adding 1M HCl. PEG_8000_ (Polyethylene glycol MW:7000-9000, Fisher Bio-Reagents, USA) was added to a final concentration of 12% (w/v), and NaCl was added to a final concentration of 0.5 M or 1.5 M. Samples were stirred at 60 rpm for 60 min (1 h) at 4°C ± 4°C according to El-Senousy et al. (2013).

### Centrifugation Following Flocculation

Flocculated samples were poured into sterile and RNase-treated (RNase Away RNA and DNA decontaminant, Molecular Bioproducts Inc.) 250 ml centrifuge bottles and centrifuged at 10,000 x*g* for 30 min at 4°C. Bottles were gently removed from the centrifuge and 10 ml of each supernatant was saved to calculate losses during the concentration step (labelled “supernatant”). The remaining supernatant was aspirated using a sterile 10-25 ml pipette directed away or opposite from the pellet location. The pellet was suspended in about 6 ml (± 1 ml) of PBS. MS2 and HAdV41 were enumerated in the suspended pellet as described above.

### Analyses

The % recovery of MS2 and HAdV41 following secondary concentration in the suspended pellet and supernatant was calculated for each test using the initial spiked concentration as the baseline. The remaining fraction of virus was considered “unaccounted”. Statistical analyses were performed with GraphPad Prism Software, Inc. v.7.0b. Two-way ANOVA analyses were performed using Tukey’s multiple comparisons test where a level of significance of *p*<0.05 was used.

## Acknowledgements

This study was funded in part by the research and development (R&D) fund of Stantec Consulting Ltd (NLM) and Ontario Centres of Excellence Voucher for Innovation and Productivity and TalentEdge grants (MH). The authors wish to thank Prof. Martha Brown and Casandra Mangroo, University of Toronto, for their advice on the propagation and endpoint dilution assay of adenovirus serotype 41. The authors wish to acknowledge Ms. Cassandra Brinovcar for assistance in processing samples.

